# Targeted CRISPR screening identifies PRMT5 as synthetic lethality combinatorial target with gemcitabine in pancreatic cancer cells

**DOI:** 10.1101/2020.05.25.112896

**Authors:** Xiaolong Wei, Jiekun Yang, Sara J. Adair, Cem Kuscu, Kyung Yong Lee, William Kane, Patrick O’hara, Denis Liu, Yusuf Mert Demirlenk, Alaa Hamdi Habieb, Ebru Yilmaz, Anindya Dutta, Todd W. Bauer, Mazhar Adli

**Affiliations:** Department of Biochemistry and Molecular Genetics, University of Virginia School of Medicine, Charlottesville, VA, USA; Department of Surgery, University of Virginia School of Medicine, Charlottesville, VA, USA; The University of Tennessee Health Science Center, Transplant Research Institute, Memphis, TN, USA; The Northwestern University, Feinberg School of Medicine, Robert Lurie Comprehensive Cancer Center, Department of Obstetrics and Gynecology, Chicago, IL, USA

## Abstract

Pancreatic ductal adenocarcinoma (PDAC) remains one of the most challenging cancer to treat. Due to the asymptomatic nature of the disease and ineffective drug treatment modalities, the survival rate of PDAC patients remains one of the lowest. The recurrent genetic alterations in PDAC are yet to be targeted; therefore, identifying effective therapeutic combinations is desperately needed. Here, we performed an *in vivo* CRISPR screening in a clinically relevant patient-derived xenograft (PDX) model system to identify synergistic drug combinations for PDAC treatment. Our approach revealed protein arginine methyltransferase gene 5 (PRMT5) as a promising druggable candidate whose inhibition creates synergistic vulnerability of PDAC cells to gemcitabine. Genetic and pharmacological inhibition results indicate that of PRMT5 depletion results in synergistic cytotoxicity with Gem due to depleted replication protein A (RPA) levels and an impaired non-homology end joining (NHEJ) DNA repair. Thus, the novel combination creates conditional lethality through the accumulation of excessive DNA damage and cell death, both *in vitro* and *in vivo*. The findings demonstrate that unbiased genetic screenings combined with a clinically relevant model system is an effective approach in identifying synthetic lethal drug combinations for cancer treatment.

**STATEMENT of SIGNIFICANCE:** Identify synergistic drug combinations for PDAC is a significant unmet need. Through CRISPR screening, we discovered and validated that PRMT5 depletion creates synergistic vulnerability of PDAC cells to gemcitabine. Mechanistically, the combination impairs DNA repair, synergistic accumulation of DNA damage and cell death *in vitro* and *in vivo*.

## INTRODUCTION

Pancreatic ductal adenocarcinoma (PDAC) is the most common and aggressive form of pancreatic cancer. It arises due to abnormal growth of exocrine ductal cells, the digestive enzyme-producing cells that compose 98% of the pancreas biomass. PDAC remains one of the deadliest of any cancer type. The main reasons for this high rate of mortality are twofold. Firstly, the disease is mostly asymptomatic until the late stages. Secondly, the current treatment strategies, mainly the chemotherapy drug combinations, are relatively ineffective. Therefore, surgery, if possible, remains the only curative therapy. However, only 15-20% of PDAC patients are eligible for surgery due to the extension of PDAC to neighboring organs. For the remaining patients, standard treatment involves radiotherapy and chemotherapy combinations. Unfortunately, current chemotherapy combinations have severe side effects due to ineffective selectivity towards the PDAC tumors. Therefore, the median survival rate is only 6 months, and more than 93% of patients die within the first 5 years(Siegel, et al. 2016). As such, despite the significant increase in the survival rates of most cancers, PDAC survival remains unchanged in the last 50 years(Kamisawa, et al. 2016), and it is projected to be the second leading cause of cancer deaths in the USA by 2030(Rahib, et al. 2014). Novel drug combinations that can result in better therapeutic value are desperately needed for PDAC treatment.

A better understanding of the significant drivers of pancreatic cancer can potentially yield novel and more effective treatment strategies. To this end, cancer genome sequencing efforts identified several recurrent genetic alterations. Among these, oncogenic *KRAS* mutations are observed in 93% of PDAC tumors(Biankin, et al. 2012). Additionally, loss of function mutations in *CDKN2A*, *TP53*, and *SMAD4* tumor suppressor genes are the main recurrent mutations. Unfortunately, none of these genetic alterations are currently targetable. The major signaling pathways downstream of oncogenic *RAS* mutations such as aberrantly active MEK-ERK pathway present themselves as promising therapeutic targets(Jones, et al. 2008). However, clinical efforts to inhibit these pathways have not been successful yet for PDAC(Little, et al. 2011; Misale, et al. 2012). Therefore, identifying effective chemotherapy combinations is a critical unmet need for PDAC treatment. However, the reliance on suboptimal *in vitro* cellular models and lack of effective unbiased approaches has hampered the ability to find effective drug combinations.

In this study, we use *in vivo* CRISPR screening to identify effective and potentially synergistic lethal drug combinations for PDAC. We specifically aimed to identify new targets whose inhibition will create conditional vulnerability with the well-established existing chemotherapy, gemcitabine (Gem). Historically, **Gem** has been the first-line chemotherapy and forms the backbone of several drug combinations for the majority of PDAC patients. The gem is a designated “essential medicine” (Robertson, et al. 2016) and has been in use since 1983. In addition to being the primary chemotherapy for PDAC, it is a critical therapy in multiple other carcinomas(Mini, et al. 2006). Although a new multidrug combination (FOLFIRINOX) slightly improves the survival of PDAC patients, due to high toxicity, only a small fraction of patients tolerate this regimen(Spadi, et al. 2016). Therefore, **Gem** remains the first-line or second-line for chemotherapy for the majority of PDAC patients. We rationalize that, given the poor 5-year survival rate of PDAC patients, finding **novel drug combinations** that will synergistically increase the therapeutic effects of Gem will have significant therapeutic value.

To this end, we utilized CRISPR/Cas9 genetic knockout (KO) screenings in clinically relevant *in vivo* model systems to find novel targets that synergize with gem. We recently performed *in vivo* and *in vitro* CRISPR KO screens for 4,000 genes to identify synthetic lethal partners of inhibitors of MEK signaling, an aberrantly active pathway due to oncogenic *KRAS* mutations in PDAC(Szlachta, et al. 2018). In this study, we focused on chromatin regulators (CR) that are essential for the survival of PDAC cells in response to Gem. We aimed to identify CRs whose inhibition creates synthetic lethality when PDAC cells are treated with Gem. We, therefore, constructed a custom sgRNA library of 8,000 sgRNAs that target ~700 epigenetic regulators and performed both *in vitro* and *in vivo* screening to identify synthetic lethal partners of **Gem**. Our approach revealed that protein arginine methyltransferase gene 5 (PRMT5) as a promising druggable candidate whose inhibition creates synergistic vulnerability of PDAC cells to Gem. At the molecular level, our findings suggest that genetic depletion or pharmacological inhibition of PRMT5 results in synergistic cytotoxicity with Gem due to depleted replication protein A (RPA) levels and an impaired non-homology end joining (NHEJ) DNA repair mechanism. Thus, the combinatorial treatment results in excessive DNA damage accumulation and subsequent cell deaths both *in vitro* and *in vivo*. These findings highlight the unbiased power of CRISPR screening in relevant clinical models to identify novel and more effective combinatorial drug targets.

## RESULTS

### *In vivo* CRISPR gene KO screening

We performed the CRISPR screening using a clinically-relevant patient-derived xenograft (PDX) model of PDAC in which a patient’s tumor is propagated *in vivo* within the pancreas of athymic nude mice(Walters, et al. 2013). The PDX366 line is established from a poorly-differentiated metastatic tumor with low stromal content and mutant for *KRAS*, *P53*, and *SMAD4* but the wild type (WT) for *P16* genes(Lindberg, et al. 2014). In our CRISPR screen (**Fig.1A**), we used an 8,031 single-guide RNA (sgRNA) library targeting 619 human genes enriched for chromatin modifiers plus 360 control sgRNAs.

**Figure 1:**
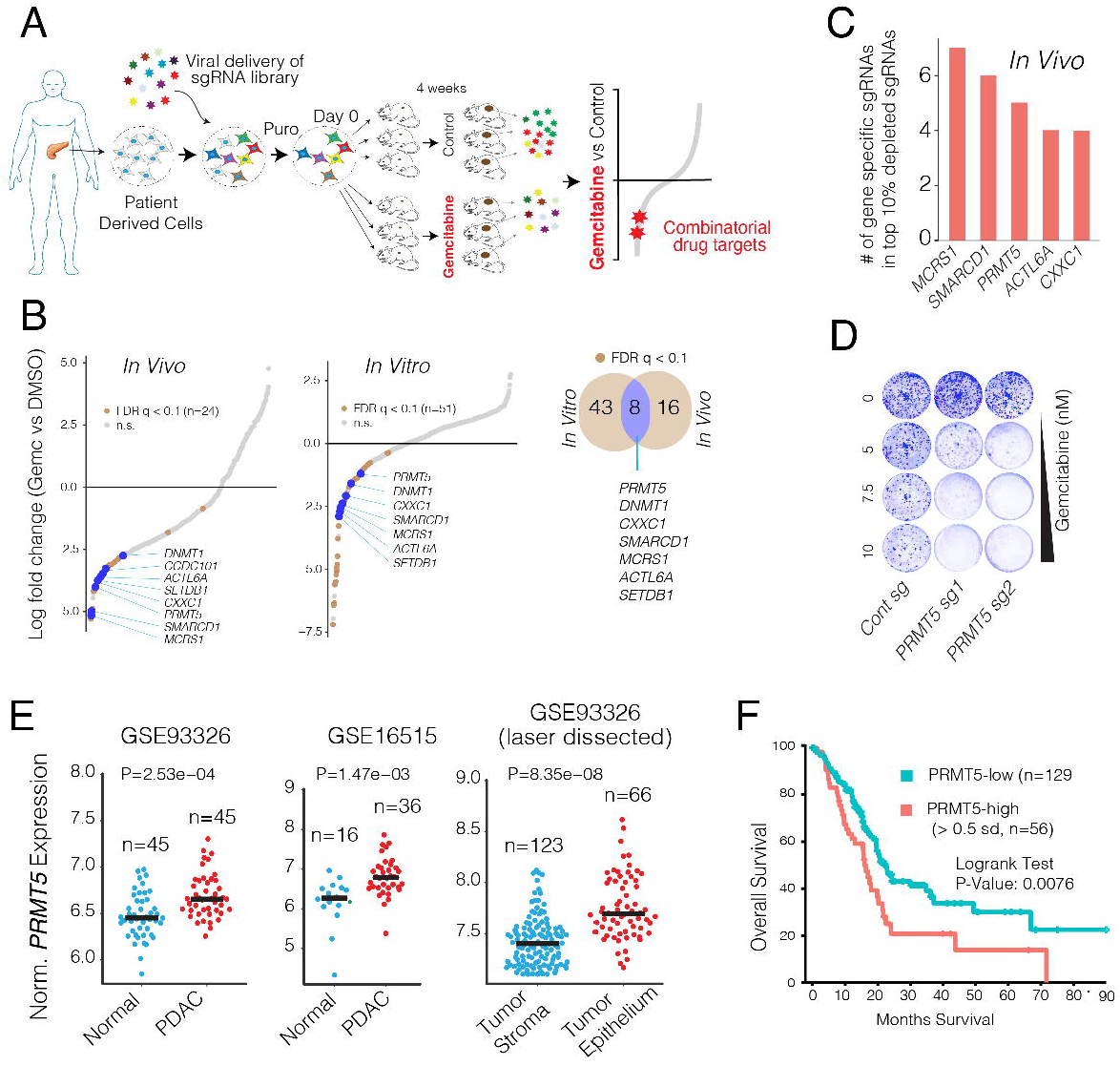
*In vivo* CRISPR screening identifies PRMT5 as a novel combinatorial target of Gem. **(A)** Schematics for *in vivo* selection screening to identify novel drug combinations. **(B)** Dot plots show gene-specific CRISPR viability scores. Significantly depleted genes (FDR< 0.1) are labeled with brown dots, whereas the genes that are depleted both *in vitro* and *in vivo* are labeled with blue dots. **(C)** Bar plots show the number of sgRNAs targeting indicated genes among the top 10% of depleted sgRNAs. **(D)** Crystal violet colony formation assays show the relative cell proliferation rates of cells expressing control and PRMT5 targeting sgRNAs in response to the indicated Gem concentrations. **(E)** Dot plots show normalized PRMT5 expression levels in normal and matched PDAC tumors. **(F)** The Kaplan-Meier plot demonstrates survival rates of PDAC patients whose tumors have high *PRMT5* expression (> 0.5 standard deviations) relative to the *PRMT5*-low patients.

To maintain sgRNA coverage, we infected ~50–100 million cells at ~0.25 multiplicity of infection (MOI). After a week of drug selection, the surviving cells were randomly divided into 9 batches, each containing ~2 million cells (~200× sgRNA coverage). Of these, one sample was harvested as “day 0,” and others were maintained in culture for *in vitro* screening or for xenograft injection into the pancreas of athymic nude mice (~2 million cells/mouse, 6 mice total). One week after injection, animals were randomized to receive either vehicle control (n = 3) or gemcitabine treatment (n = 3) for 4 weeks as described in Materials and Methods and depicted in **Fig. 1A**. The relative abundance of each sgRNA was assessed by targeted amplification and deep sequencing of tumor genomic DNA. Data analysis was performed using MAGeCK (Li et al., 2014) and R. In parallel, we also performed *in vitro* screening, in which cultured cells were exposed to control dimethyl sulfoxide (DMSO) or 20% inhibitory concentration (IC_20_) doses of Gem every 3 days for 4 weeks.

The sgRNA read count distribution analyses of the day 0 sample (> 99.9% coverage) demonstrated that the sgRNAs in our library were evenly represented with a Gini index of 0.07 (~0.1 is suggested for initial-state samples(Koike-Yusa, et al. 2014), **Supplementary Fig. 1**). Contrary to the day 0 samples, 86%, and 81% of the sgRNAs were detectable in control *in vitro* and *in vivo* samples after a month of selection, respectively. Assuming that the ~15% depletion was due to the functional roles of the target genes, the analyses suggested that ~95% of cells containing sgRNAs contributed to *in vivo* tumor formation (**Supplementary Fig. 2**). Notably, only 20 and 12 genes had 2 or fewer sgRNAs in the *in vitro* and *in vivo* control samples, and 7 genes were overlapped **(Supplementary Fig. 3)**. The 7 genes include essential DNA repair genes like *CHEK1*, *MSH2*, and *RAD21* **(Supplementary Fig. 3)**. One of the *in vivo* Gem treated tumors had substantially more sgRNAs depleted compared to the other two replicates. Reasoning that this tumor responded to gemcitabine at a higher than expected rate, we excluded it from the downstream analysis (**Supplementary Fig. 2**). Since the non-genomic targeting control sgRNAs were well-represented in all of the samples, they were used to profile the null distribution of Robust Rank Aggregation (RRA) scores when calculating the P values(Li, et al. 2014) (**Supplementary Fig. 4**). Negative selection RRA scores identified genes that were consistently depleted when compared to day 0 samples, indicating that these genes are critical for the survival of the PDX cell line (**Supplementary Fig. 5**). To check this, we compared these set of genes with known essential fitness functions. Critically, more than half of the 104 critical survival genes that we identified from the *in vitro* and *in vivo* samples overlapped with the previously identified essential gene list from five independent cell lines(Hart, et al. 2015). It is also notable that nearly 1/3 of essential fitness genes we identified are *in vitro* or *in vivo* specific, indicating their differential essentiality for different growth conditions. Reasoning the limited therapeutic index of targeting these essential genes, we excluded them from the candidate genes that showed synthetic lethality with Gem.

### Identifying genes whose depletion results in synthetic lethality with gemcitabine

We aimed to identify genes that could be therapeutically targeted to synergistically boost the therapeutic effect of Gem. We, therefore, prioritized our CRISPR screening hits based on three criteria. Firstly, the gene must have been significantly depleted both *in vitro* and *in vivo*. We also included the *in vitro* screening data so that we could robustly validate the screening hits using *in vitro* assays. Secondly, the potential hit must have been “druggable,” i.e., have an existing small molecule inhibitor. And finally, targeting the CRISPR hit should have a high potential for strong therapeutic value. Among these three criteria, the latter one is more ambiguous. To this end, we focused on genes whose high expression has strong negative prognostic value for PDAC patients.

This **primary screening (Fig. 1A)** identified *MCRS1, SMARCD1, PRMT5, CXXC1, SETDB1, ACTL6A*, and *DNMT1* as significant hits whose depletion was potentially lethal with Gem (Fig. 1B **and** C). We then performed a validation screening with additional sgRNAs for each of these genes using the PDX366 cell line. *PRMT5* scored as the top hit whose depletion synergistically increased Gem cytotoxicity (**Fig. 1D**). PRMT5 is the primary type II PRMT that is responsible for the majority of symmetric dimethylation (SDMA) on the arginine residues of its targets, which include various histone proteins as well as transcription factors(Blanc and Richard 2017). PRMT5 is implicated in diverse functions, including genome organization, transcription, cell cycle, and spliceosome assembly(Stopa, et al. 2015). The role of PRMT5 in the proliferation of cancer cells is increasingly appreciated(Clarke, et al. 2017; Hamard, et al. 2018; Jansson, et al. 2008; Zheng, et al. 2013). Importantly, PRMT5 is a druggable protein with several selective inhibitors available and many of which are currently tested in clinical trials (e.g., NCT03573310 and NCT03854227). However, its significance in PDAC progression or its potential as a combinatorial therapeutic target in PDAC cells has not been explored.

We, therefore, investigated the potential role of PRMT5 in PDAC progression and aimed to assess whether PRMT5 inhibition has a potential therapeutic value for PDAC. To this end, we initially analyzed whether PRMT5 expression is differentially regulated in PDAC tumors and has a prognostic value for the survival of patients. The gene expression analysis of multiple independently generated datasets such as normal-matched PDAC tumor (GSE28735)(Zhang, et al. 2013), tumor-adjacent normal vs PDAC tumor (GSE16515)(Pei, et al. 2009), and laser microdissected PDAC tumor cells vs. adjacent stromal cells (GSE93326)(Renz, et al. 2018) shows that *PRMT5* mRNA expression is significantly upregulated in PDAC cancer cells compared to normal stromal cells (**Fig. 1E)**. Most critically, the analysis of TCGA PDAC patient data shows that tumors with high PRMT5 expression result in significantly shorter overall patient survival **(Fig. 1F**), indicating that PRMT5 is a critical player in PDAC progression or therapy response, and thus a promising therapeutic target.

To better study the role of PRMT5 in the cellular response to Gem, we generated multiple single KO clones in two additional pancreatic cancer cell lines (mPanc96 and PANC-1) **(Fig. 2A**). Notably, despite screening for ~100 single clones, we seldom observed full depletion of PRMT5 at the protein level, especially in PANC-1 cells. However, the clones with even partial PRMT5 depletion were nearly an order of magnitude more sensitive to Gem compared to wild-type (WT) clones (4-5 uM vs. ~50 uM IC_50_) as measured by cell viability assay **(Fig. 2B)**. In line with these, longer-term crystal-violet colony formation assays also demonstrated that the PRMT5 KO clones were significantly more sensitive to Gem compared to WT cells **(Fig. 2C)**.

**Figure 2:**
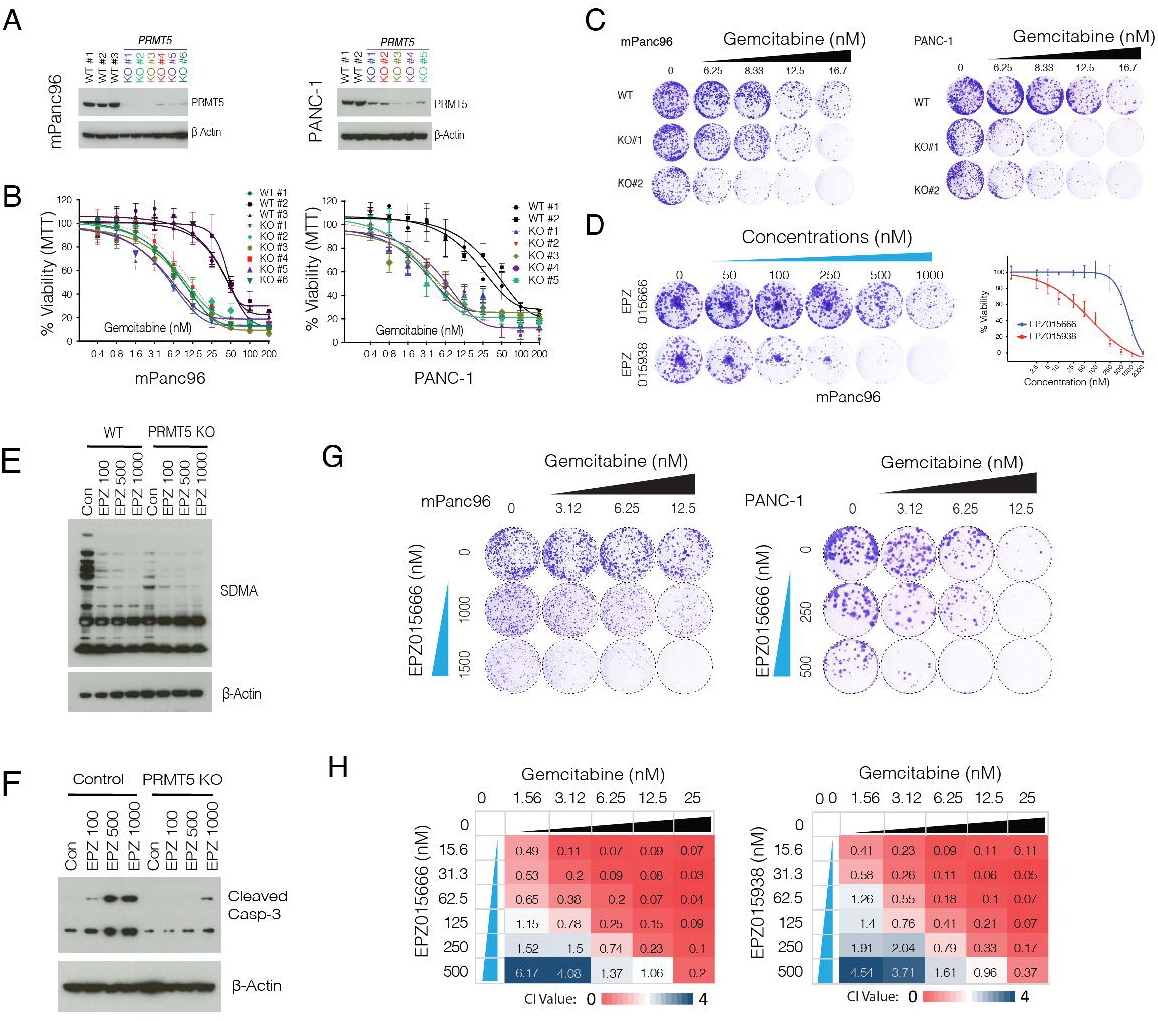
*PRMT5* depleted cells are hypersensitive to Gem. **(A)** Western blot shows PRMT5 protein levels in WT cells, as well as single-cell, expanded clones expressing PRMT5 targeting sgRNAs. The beta-actin level is shown as a loading control. **(B)** The line graph shows % viability of WT and *PRMT5* KO PDAC cells in response to increasing doses of Gem. **(C)** Crystal violet colony formation assay shows the overall survival of indicated WT and *PRMT5* KO cells. **(D)** Crystal violet colony formation assay shows relative growth inhibition activity of two separate PRMT5 inhibitors. **(E)** Western blot result shows relative levels of symmetric dimethylation of arginine (SDMA) in WT cells and PRMT5 KO cells treated with increasing concentration (nM) of the indicated PRMT5 inhibitor. **(F)** Western blot result shows a relative rate of Caspase-3 cleavage in WT and PRMT5 KO cells treated with increasing doses (nM) of PRMT5 inhibitor. **(G)** Crystal violet colony formation assay shows relative survival and proliferation rates of mPanc96 (left) and PANC-1 cells (right) treated with various combinatorial doses of Gem and two separate PRMT5 inhibitors. **(H)** Heatmaps show the Combination Index (CI) values across multiple combinatorial doses in PDX 366T cells. CI<1 indicates synergism.

To further corroborate these genetic depletion results, we tested two separate small molecule pharmacological inhibitors (EPZ015666 & EPZ015938) that specifically target PRMT5. When tested as a single agent, EPZ015938 had substantially more growth inhibition activity on colony formation **(Fig. 2D)**. As anticipated, EPZ015666 treatment significantly inhibits global SDMA (**Fig. 2E**). Furthermore, the inhibitor is specific towards PRMT5 as it results in significant apoptosis (caspase-3 cleavage) selectively in WT cells but not in PRMT KO cells (**Fig. 2F**). These inhibitors significantly potentiated Gem growth inhibition activity at multiple dose combinations as measured by long-term colony formation assay **(Fig. 2G)**. To better assess whether PRMT5 inhibitors are synergistic with Gem, we calculated the **Combination Index (CI)** values for each dose combination(Chou and Talaly 1977). The CI < 1 indicates synergy between two drugs, whereas CI ≈ 1 is additive, and CI ≥ 1.2 suggests an antagonistic effect. Importantly, of the 24 dose combinations for two separate inhibitors, we observed robust synergistic activity for ~80% of EPZ015666 + Gem and ~78% of EPZ015938 + Gem dose combinations **(Fig. 2H)**.

### Understanding pathways underlying PRMT5 depletion-mediated vulnerability to Gem

Encouraged by the genetic depletion and pharmacological inhibition studies, we then aimed to understand the molecular mechanism of conditional sensitivity to PRMT5 depleted cells to Gem and assess the therapeutic value of this combination *in vivo*. At the chemical level, Gem is composed of difluoro-deoxycytidine (**dFdC**). Mechanistically, it exerts its biological effects by inducing replication stress in fast-dividing cancer cells. Once taken up by the cells, **dFdC** is converted into dFdC-diphosphate (dFdCDP) and dFdC-triphosphate (dFdCTP). dFdCTP incorporates into DNA as a cytosine analog and blocks DNA synthesis due to strand termination. Additionally, dFdCDP also inhibits the ribonucleotide reductase enzyme, thereby resulting in depletion of the dNTP pool necessary for DNA synthesis.

Since PRMT5 is a major transcriptional regulator, we initially investigated whether PRMT5 depleted cells had a differential transcriptional response to Gem. We therefore, comparatively analyzed the transcriptional responses of *PRMT5* WT and KO cells to Gem in two independent PDAC cancer cell lines. Critically, the KO cells responded to Gem by differentially regulating a much larger number of genes. For example, while only 21 genes (9 up, 12 down) in WT PANC-1 cells and 512 genes (252 up, 260) in mPanc96 WT cells were significantly altered, 1,589 genes (918 up, 680 down) in PANC-1 KO cells and 1,385 genes (920 up, 465 down) in the mPanc96 KO cells were significantly altered in response to treatment with IC_30_ Gem for 24 hours **(Fig. 3A)**. These results suggested that physiological levels of PRMT5 are required to buffer global transcriptional response to Gem. Comparative gene set enrichment analysis demonstrated that genes implicated in cell cycle, and DNA repair pathways were aberrantly active in the KO cells when treated with Gem **(Fig. 3B)**. Gene sets identifying cell cycle-related genes such as G2/M checkpoints, and E2F and MYC targets were all more strongly upregulated in Gem treated KO cells compared to WT cells. Furthermore, genes involved in DNA repair were among the most highly differentially regulated genes when the KO cells were treated with Gem **(Fig. 3B)**.

**Figure 3:**
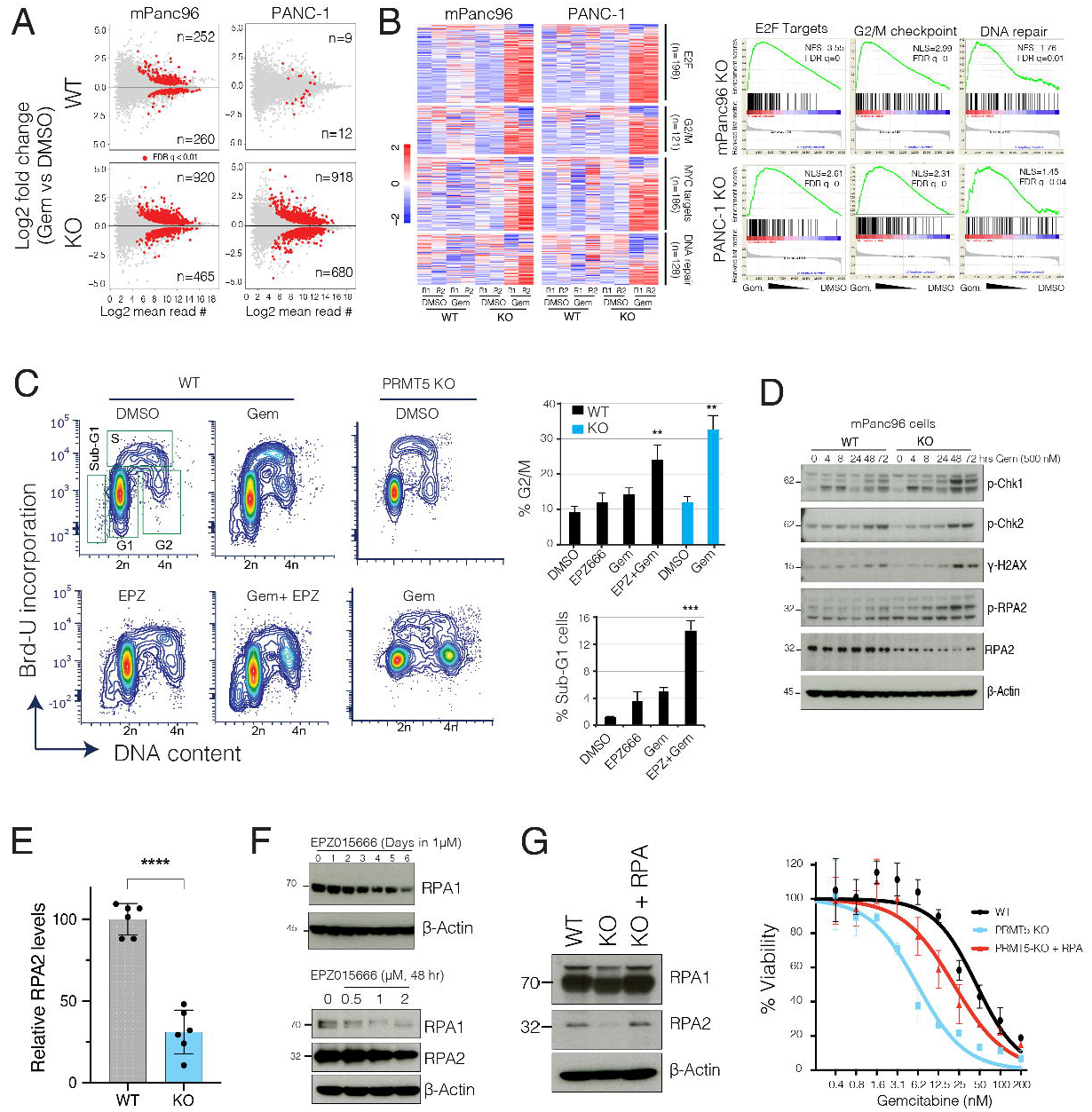
*PMRT5* depletion results in the aberrant transcriptional program of cell cycle and DNA repair genes in response to Gem. **(A)** The MA plots (log fold change vs. log mean expression of each gene) show the number of differentially regulated genes in WT and PRMT5 KO cells due to Gem treatment. **(B)** Heatmaps and Gene Set Enrichment Analysis (GSEA) show relative levels of expression changes in genes involved in indicated cellular processes. **(C)** Flow cytometry cell cycle analysis (DNA content vs. BrdU incorporation) of control vs. Gem (250 nM) treated WT, PRMT5 KO, or PRMT5 inhibitor EPZ015666 (500nM) treated cells. The bar plot shows the percent of cells at the indicated cell cycle stage. ** & *** indicates p values less than 0.01 and 0.001, respectively. **(D)** Western blots show relative levels of phosphorylated or total levels of indicated proteins. **(E)** Bar plots of RPA2 protein levels quantified from western blots. **(F)** Western blots show levels of RPA1 and RPA2 proteins in PDAC cells treated with various time and doses of the indicated PRMT5 inhibitor. **(G)** Western blots show relative levels of RPA1 and RPA2 protein levels in WT, PRMT5 KO, and PRMT5 KO cells expressing RPA cDNA. The line plots show the relative viability of indicated cells in response to increasing doses of Gem.

These results support the hypothesis that *PRMT5* depletion-mediated conditional vulnerability to Gem was partially due to the aberrant regulation of cell cycle and DNA repair pathways. To test this hypothesis, we set out several molecular assays to study the mechanism of *PRMT5* depletion mediated aberrant cell cycle and DNA repair programs. The analysis of cell cycle position through BrdU incorporation showed that Gem treatment of WT cells resulted in a partial delay in cell cycle with a substantial accumulation of cells in S-phase and partial increases in G2/M cells **(Fig. 3C)**. On the other hand, the combination of Gem and PRMT5 inhibitor resulted in a significant accumulation of G2/M cells and sub-G1 dead cells **(Fig. 3C)**. In line with the pharmacological inhibition of PRMT5, Gem treatment resulted in a significantly higher number of G2/M cells in the *PRMT5* KO cells compared to WT cells.

### PRMT5 depletion results in RPA exhaustion

Coordinated activation of cell cycle checkpoints and cell cycle arrest is one of the primary mechanisms that enable cells sufficient time to repair DNA against external cues(Curtin 2012). The robust arrest of cells at the G2/M cell cycle led us to study the activation of checkpoints further. The S and G2/M cell cycle arrest results from DNA damage that mediates activation of ATR-Chk1-Cdc25C(Bartek, et al. 2004; Blackford and Jackson 2017; Marechal and Zou 2013). We, therefore, performed time-course experiments to study whether Gem treatment resulted in differential activation of DNA damage and cell cycle checkpoints in the KO cells. Notably, we detected sustained and stronger phospho-Chk1 (a marker of DNA damage, S and/or G2/M arrest), gamma-H2AX (a marker of DNA damage) as well as phospho-RPA2 (a marker of DNA damage and replication stress) in the KO cells compared to WT cells (**Fig. 3D**).

This analysis also revealed something unexpected to us. Although we observed a strong induction of phospho-RPA2 in the KO cells, the total levels of RPA2 were substantially lower in the KO cells (**Fig. 3D**). Further quantitative analyses suggested that the depletion of *PRMT5* resulted in a significant reduction in RPA2 protein levels (p<0.0001). We observe a ~60-70% reduction in RPA2 levels in the *PRMT5* KO cells compared to WT cells **(Fig. 3E)**. These findings led us to investigate whether the depletion of RPA was due to enzymatic activity of PRMT5. Critically, both time-course, as well as dose-escalation experiments, showed that depletion of PRMT5 activity through small molecule inhibitors resulted in RPA2 depletion **(Fig. 3F)**.

It should be noted that RPA2 is one of the three subunits of the RPA complex, which is viewed as the guardian of the genome(Liu and Huang 2016; Wu, et al. 2016), because it binds and protects any single-stranded DNA that forms during DNA replication, transcription and repair pathways. The cytotoxicity of Gem in fast-dividing cancer cells is mostly due to the creation of **replication stress** (**RS**) by blocking DNA synthesis and diminishing the dNTP pool by inhibiting ribonucleotide reductase enzyme. During replication stress, RPA becomes essential to protect single-strand DNA (ssDNA) at the stalled replication forks. Critically, overall RPA levels are crucial determinants as to whether cells can resolve the stalled forks. In low RPA conditions, the replication stress leads to “**replication catastrophe**,” where chromosomes chatter with thousands of double-strand breaks (DSB)(Toledo, et al. 2013). These findings lead to the “**RPA exhaustion**” hypothesis, which states that when RPA is not sufficient, cells can’t survive the replication stress, and the stalled replication forks **collapse**, which results in the **breakage** of forks, and ultimately replication catastrophe (Toledo, et al. 2017).

Our results so far support the hypothesis that *PRMT5* depletion results in “RPA exhaustion,” and therefore, cells are not able to cope with Gem-mediated DNA damage. To test this hypothesis, we aimed to replenish the RPA complex to see if it could rescue the *PRMT5* depletion phenotype. Since RPA works as a tri-partite complex where each subunit is needed at an equimolar ratio, we exogenously provided cells with a vector that expresses all three subunits(Toledo, et al. 2013). Importantly, replenishing the RPA complex in *PRMT5* KO cells to near equal levels to WT cells (**Fig. 3G**) results in significant resistance to Gem in *PRMT5* KO cells. These results suggest that, at the molecular level, the *PRMT5* depletion-mediated Gem-sensitivity phenotype is, in part, due to exhaustion of the RPA complex.

### PRMT5 depletion results in excessive DNA damage accumulation

Our expression analysis also highlighted that genes involved in DNA repair pathways were aberrantly regulated when *PRMT5* KO cells were treated with Gem. Furthermore, the above results suggested that due to RPA exhaustion, PRMT5 depleted cells are not able to resolve a stalled replication fork, which may result in the collapse of the fork and accumulation of DNA DSB. We, therefore, utilized two independent molecular assays to detect and quantify Gem-induced DNA damage in control and *PRMT5* depleted cells. Initially, we used immunofluorescence (IF) assays to detect the phosphorylated H2AX (γ-H2AX), which is a modified histone variant deposited into an around megabase chromatin region around DSB. Critically, strong γ-H2AX foci can be detected as early as 4 hours post Gem treatment in the KO cells. On the other hand, it took 48-72 hours to detect similar levels of γ-H2AX foci in WT cells using the same concentration of Gem **(Fig. 4A)**. Quantitative analysis of γ-H2AX foci formation levels over a period of 72 hours showed that *PRMT5* KO cells consistently had significantly higher levels of γ-H2AX, indicating higher levels of DNA DSB due to Gem treatment **(Fig. 4A**, lower panel, bar plots). In line with the genetic depletion of *PRMT5*, pharmacological inhibition of PRMT5 with two separate small molecule inhibitors also demonstrated that depletion of PRMT5 activity resulted in significant accumulation of DNA DSB, as detected by levels of γ-H2AX foci formation (**Fig. 4B**). In addition to γ-H2AX foci formation, we also measured the level of DNA strand breaks through a comet assay, which measures the overall levels of DNA damage, as done through single-cell gel electrophoresis. As the frequency of DNA breaks increases, so does the fraction of the DNA extending towards the anode, forming the comet tail. The length of the tail is an indication of levels of fragmented DNA in individual cells. In line with the γ-H2AX IF results, we observed a significantly longer comet tail when *PRMT5* KO cells or PRMT5-inhibited cells were treated with Gem compared to WT and control-treated cells, respectively (Fig. 4C **and** D).

**Figure 4:**
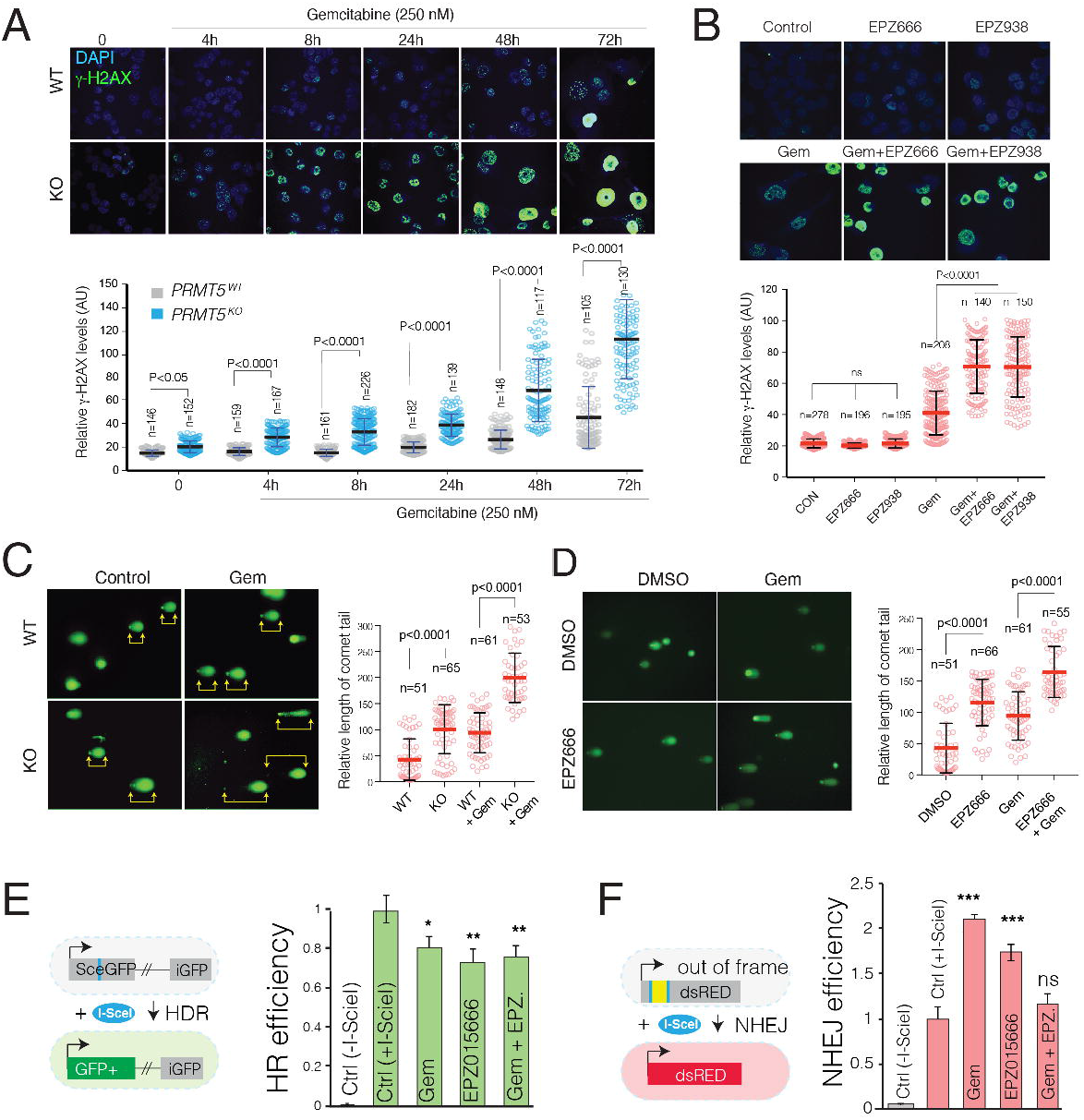
*PRMT5* depletion results in impaired DNA repair and excessive DNA damage accumulation in the PDAC cells treated with Gem. **(A-B)** Immunofluorescent (IF) images of γ-H2AX relative levels in WT, *PRMT5* KO, and PRMT5 inhibitor EPZ015666 or 938 (500nM) treated WT mPanc96 cells in response to Gem treatment (250 nM) for indicated times (Upper panels). The dot plot in the lower shows quantified IF γ-H2AX levels at indicated number of single cells. N indicates the number of cells quantified. **(C-D)** IF images of Comet assay indicate, levels of overall DNA strand breaks in WT, PRMT5 KO, and EPZ015666 (500nM) treated WT mPanc96 cells in response to Gem treatment (250 nM) (Upper panel). The lower panels show individual cell level quantified length of the comet tail in the indicated number of cells. **(E-F)** The bar plot shows results of I-SceI endonuclease-based genetic reporter assays indicating relative repair efficiency of DNA strand breaks through homology-directed repair (HDR) (E) or non-homology end joining (NHEJ) pathways in HeLa cells treated with Gem (250 nM) and/or EPZ015666 (500 nM) (F).

### PRMT5 depletion results in impaired NHEJ

Depending on the time and kind of DNA damage, DSB is repaired through either precise homology-directed DNA repair (HDR) or error-prone NHEJ(Curtin 2012). NHEJ is active throughout the cell cycle, whereas HDR is restricted to the late S and G2 phases of growing cells(Panier and Boulton 2014). Our differential gene expression results, as well as RPA exhaustion findings, led us to investigate whether excessive DNA damage accumulation was, in part, due to impaired DNA repair activity. To this end, we used **I-SceI** endonuclease-based genetic reporters where relative efficiency of DSB repair by either pathway could be robustly quantified. In the NHEJ reporter assay, the dsRED contains a “stuffer sequence” flanked by two I-Sce recognition sites, which puts dsRED out of frame. On the other hand, for the HDR GFP-reporter system, the construct contains two defective GFP genes, the first one contains an I-SceI site. In both cases, the engineered HeLa cells are dsRed (−) or GFP(−), respectively. However, exogenous expression of I-SceI leads to a DSB repair and creation of either dsRED+ or GFP+ cells, which can be quantified to assess relative NHEJ or HDR repair efficiencies by quantifying the percentage of dsRED+ or GFP + cells upon I-SceI expression(Golding, et al. 2009; Gunn and Stark 2012). Our results show that either Gem and PRMT5 inhibitor treatment significantly inhibits the HDR activity **(Fig. 4E)**. Notably, the combination treatment did not result in any further reduction in HDR activity, suggesting that the reduced HDR activity did not explain the observed synergistic accumulation of DNA damage. We then assessed whether PRMT5 inhibitor alone or in combination with Gem results in differential NHEJ activity. Importantly, unlike HDR activity, we observed a significant increase in NHEJ activity when cells were treated with either Gem or PRMT5 inhibitor. Surprisingly, when cells were treated with the Gem plus PRMT5 inhibitor combination, there was no significant change in NHEJ activity. This finding supports a hypothesis that reduced HDR repair due to a single Gem or PRMT5 in treatments is compensated by an increase in NHEJ. However, the combination treatment was not able to increase the NHEJ repair pathway and thus cannot compensate for the reduced HDR activity.

### The combinatorial treatment results in synergistic tumor growth inhibition *in vivo*

Next, we investigated to see if the combination would result in synergistic growth inhibition of PDAC tumors. To this end, we explored both genetic depletion and pharmacological inhibition of PRMT5 in a xenograft model of PDAC. Initially, we tested whether tumors formed by WT and *PRMT5* KO cells were differentially sensitive to Gem treatment. To be able to better compare the tumors from these two genetic backgrounds, we injected 5×10^5^ WT mPanc96 cells in the left flank and the same number of the *PRMT5* KO cells in the right flank of the same mouse. This strategy enabled us to compare the two tumors grown in the same mouse. After one week of tumor formation, the mice were randomly divided into three groups where one received control, and the other two received two separate Gem doses (50 mg/kg or 100 mg/kg). Notably, the *PRMT5* KO cells were able to form tumors. However, these tumors *were* slightly smaller than the tumors formed by WT cells. The Gem treatments did result in a notable reduction of WT tumors. However, the most significant reduction of tumor volumes was observed in the *PRMT5* KO tumors treated with Gem (Fig. 5A **and** B). Starting from the fifth treatment (day 20 of tumor formation), the Gem treated *PRMT5* KO tumors were significantly smaller than their WT counterparts or the untreated KO tumors.

**Figure 5:**
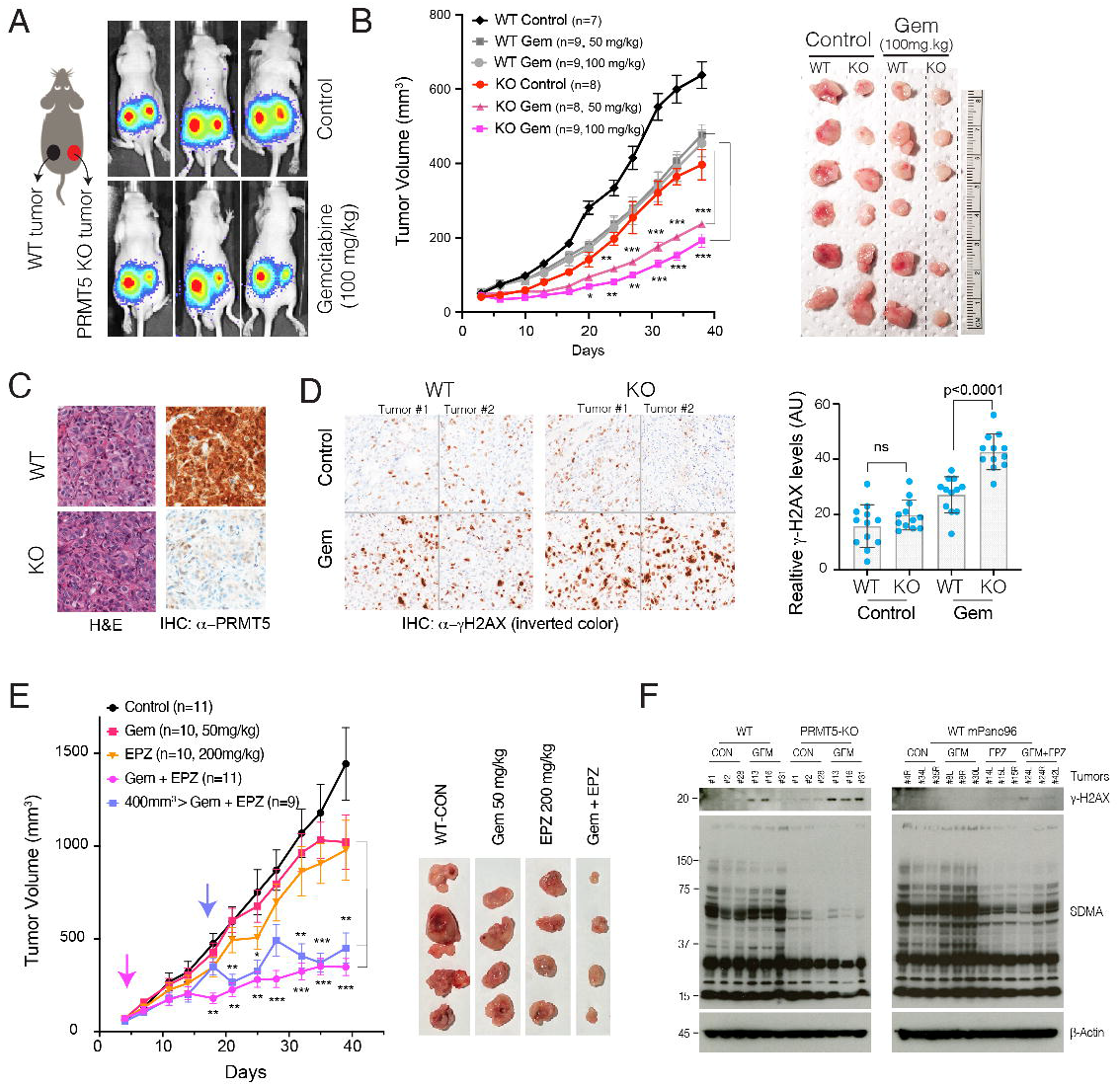
Genetic depletion or pharmacological inhibition of PRMT5 together results in synergistic tumor growth inhibition with Gem. **(A)** Schematics show the experimental strategy where WT and *PRMT5* KO mPanc96 cells are xenografted in the left and right side of the mice, respectively. The bioluminescence imaging results show relative levels of WT and PRMT5 depleted tumors in control and Gem treated mice. **(B)** The line plot shows caliper-measured relative tumor volumes over time in WT and PRMT5 depleted tumors treated with control and two separate Gem doses. The images show extracted tumors at the end of the experiments. **(C)** Hematoxylin and eosin (H&E) and immunohistochemistry (IHC) stainings respectively show tumor architecture and relative levels of PRMT5 protein in tumors originated from WT and *PRMT5* KO cells. **(D)** IHC images and bar plots show relative levels of DNA damage (γ-H2AX staining) in WT and *PRMT5* depleted tumors treated with control vehicle or Gem. **(E)** The line plots show caliper measured relative tumor volumes in vehicle control, single agent or combinatorial Gem, and PRMT5 inhibitor (EPZ015666) treated mice. Pink and blue arrows indicate treatment start times for respective modalities. The images show extracted tumors at the end of the experiment. **(F)** Western blots indicate relative levels of DNA damage (γ-H2AX) and SDMA in multiple different tumor tissues receiving indicated treatments.

We also extracted tumors to analyze their morphology and molecular structure through H&E and IHC for selected markers. The H&E staining of WT and *PRMT5* KO tumors demonstrated comparable cellular architecture **(Fig. 5C)**. The IHC staining confirmed that the tumors from *PRMT5* KO cells did not express PRMT5 protein **(Fig. 5C)**. We then performed IHC to investigate whether the Gem treatment resulted in greater DNA damage *in vivo* than was seen in *in vitro* experiments. Consistent with the *in vitro* experiments, we observed significantly more γ-H2AX staining in the *PRMT5* KO tumors when treated with Gem **(Fig. 5D)**, indicating that Gem resulted in a significantly higher amount of DNA damage in the PRMT5 KO tumors than in the WT tumors.

We next performed *in vivo* xenograft experiments to assess whether pharmacological inhibition of PRMT5 would result in a synergistic reduction of PDAC tumors in vivo when combined with Gem. To this end, we designed two separate strategies: early treatment and delayed treatment. One set of tumors were treated with control, single-agent drugs or combination of Gem and PRMT5 inhibitor as soon as the tumors reached ~100 mm^3^. Importantly, as soon as a week after treatment, we observed a statistically significant reduction in tumor volume only in the tumors receiving the combination treatment. It is notable that the therapeutic effect of PRMT5 inhibitor plus Gem is much stronger than the Gem or PRMT5 inhibitor treatment alone (**Fig. 5E**). Importantly, we also allowed a set of tumors to grow to significant sizes (~400 mm^3^) before starting the combination treatment. Interestingly, we observed a notable decrease in tumor volume after just one cycle of combined treatment (**Fig. 5E**). These tumors started to grow for the next couple of treatment cycles but then stopped growing and remained significantly smaller in volume compared to the control or single-agent treated tumors. In the end, these tumors were almost indistinguishable from the tumors that received combination treatment from the beginning.

To assess whether the combination treatment abolished PRMT5 function *in vivo*, we evaluated the overall levels of the γ-H2AX and the asymmetric arginine dimethylation (SDMA) in WT tumors, *PRMT5* KO tumors as well as single-agent and combination-treated WT tumors. As anticipated, the *PMRT5* KO tumors had significantly lower SDMA levels. In line with genetically depleted tumors, the EPZ015666 treated tumors had lower overall levels of SDMA, indicating that the inhibitor doses that we used resulted in substantial inhibition of PRMT5 function in the tumors *in vivo* (**Fig. 5G**). However, the γ-H2AX is significantly activated in either the GEM-treated PRMT5 KO tumors or the combination-treated tumors (Fig. 5G), suggesting that activation of the γ-H2AX may be due to the low level of SDMA.

## DISCUSSION

Our main objective in this study was to utilize an *in vivo* CRISPR screen to identify novel drug targets that could synergistically increase the therapeutic efficacy of existing chemotherapy in PDAC, which remains one of the deadliest despite extensive efforts. It comprises 95% of pancreatic cancers, and the 5-year survival rate is less than 8%. Rationalizing that the nucleoside analog Gem is the most widely used chemotherapy agent and the backbone of other combinatorial therapies(Spadi, et al. 2016), we specifically focused on identifying novel combinatorial targets that may result in synthetic lethality with Gem. To achieve this, we employed the CRISPR technology with a library of sgRNAs to screen chromatin regulators whose depletion may create conditional lethality with Gem. The screening was performed in an orthotopic PDX model where a tumor is growing in the pancreas of the mice. In this screening, we used a custom sgRNA library targeting a ~700 genes implicated in epigenetic control of chromatin regulation. The initial screening and subsequent validation experiments demonstrated that PRMT5 inhibition resulted in the synergistic vulnerability of PDAC cells to Gem.

The role of PRMT5 as a critical driver of cancer progression is an emerging topic. More importantly, PRMT5 is a druggable protein. There are several ongoing clinical trials of PRMT5 inhibitors for multiple advanced-stage cancers (NCT03573310, NCT03854227). Although PDAC patients are currently not recruited for these trials, our findings presented here show that PRMT5 is a significant therapeutic target in PDAC. Firstly, PRMT5 expression is significantly induced in PDAC tumors. Secondly, higher PRMT5 expression in PDAC tumors is significantly associated with poor patient survival. These findings already indicate that inhibiting PRMT5 will positively impact patient survival. Notably, our screening results indicate that PRMT5 inhibition is synthetic lethal with Gem. Since PDAC cells express higher levels of PRMT5, these findings strongly suggest that combinatorial PRMT5 inhibition with Gem will have strong therapeutic effects.

Recently, several complementary reports demonstrated that combining PRMT1 inhibitors with PRMT5 inhibitors resulted in synergistic anti-tumor activity(Fedoriw, et al. 2019; Fong, et al. 2019). Although PRMT1 did not score as one of the top hits in our screening, it is tempting to speculate that the triple combination of inhibitors targeting PRMT1 and PRMT5 with Gem may result in far greater synergistic cytotoxicity towards PDAC cancer cells. In such a combinatorial strategy, achieving a high therapeutic index is the ultimate goal in cancer treatment. Notably, the PRMT5 inhibitor dose that we use achieved significant and synergistic cytotoxicity with Gem and is an order of magnitude less than what has been shown to have a cytotoxic effect in hematopoietic cancers (250 nM vs 10 μM)(Hamard, et al. 2018). These results indicate that combinatorial strategies may have a high therapeutic index.

How PRMT5 inhibition results in synergistic cytotoxicity with Gem is yet to be understood entirely. Our results indicate that PRMT5 depleted cells are not able to tolerate the replication stress elicited by the Gem treatment. These findings are in line with the reports that highlight PRMT5 as a critical regulator of DNA damage response(Clarke, et al. 2017; Hamard, et al. 2018). At the molecular level, our findings support the hypothesis that PRMT5 depletion results in a substantial “RPA exhaustion” model. This model highlights that RPA levels are critical determinants of whether a stalled replication fork resolve or lead to what is called “replication catasrophe(Toledo, et al. 2013; Toledo, et al. 2017)”, a molecular process that results in chromosome chattering due to excessive DNA DSB. In addition to RPA depletion, our findings indicate that PRMT5 inhibition, in combination with gem, results in defective DNA repair by NHEJ.

In conclusion, our findings indicate the power of unbiased *in vivo* CRISPR screening in a clinically relevant model system. One limitation of *in vivo* screening is the limited number of target genes that can be screened. This limitation emerges due to the maximum number of cells that can be injected into the relevant anatomical site in the *in vivo* model. Since the complexity of the sgRNA library needs to be preserved at ~200× during all experimental steps, screening with a library targeting the whole genome is currently not feasible with the CRISPR/Cas9 system. However, alternative CRISPR systems, such as the CRISPR/AsCpf1 system, can be exploited to construct significantly smaller libraries(Liu, et al. 2019), which may potentially enable genome level screening *in vivo*.

## Materials and Methods

### In vitro cell culture

Human PDX366 (Patient-derived pancreatic tumor cells), mPanc96, and PANC-1 pancreatic carcinoma cells were cultured in RPMI1640 medium supplemented with 10% fetal bovine serum (FBS) and 1% streptomycin/penicillin. Cells were treated with gemcitabine (GEMZAR; Eli Lilly) and/or either EPZ015666 (GSK3235025; SelleckChem) or EPZ015938 (GSK3326595; ChemieTek).

### Generation of CRISPR sgRNA library pool and viral infection

PDX366 cell line was produced from pancreatic patients, as described earlier(Walters, et al. 2013). WT Cas9 expressing lentivirus was generated in HEK293T cell line by co-transfection of WT Cas9 (modified from GeCKO plasmid by removing gRNA), psPAX and pMD2.6 plasmid with 5:4:1 ratio. Ten μg total DNA was used in the presence of 30 μl of Fugene6 reagent in a 10-cm plate dish that had 70 % confluency. PDX366 cell line was infected with this lentivirus for one day and then treated with 0.5 μg/ml puromycin for four days. The nuclear sgRNA libraries were kind gifts from Dr. Sabatini lab (MIT) (*Wang et al.*(Wang, et al. 2014); Addgene Catalogue # 51047). The libraries were amplified using the published protocol at Addgene. (http://www.addgene.org/static/data/08/61/acb3ad96-8db6-11e3-8f62-000c298a5150.pdf). The library pool targets 619 epigenetic regulators with ~ ten sgRNA/gene. 360 nongenomic targeting control sgRNAs are included in the library. The sgRNA library expressing viruses was generated in 2× 15 cm plates by using a total of 20 μg DNA and the condition mentioned above. Serial dilutions of a virus were used to find the MOI of ~0.25 after selection with five μg/ml blasticidin for four days. Cells were harvested from 12×15 cm plates to get at least 200× fold coverage (~2 million cells per sample) for the *in vitro* and *in vivo* (orthotopic injection into mouse pancreas) screening.

### *In vivo* CRISPR screening in an orthotopic patient-derived xenograft (PDX) model of PDAC

6-7 week-old athymic nude mice (Envigo, Indianapolis, IN) were used for in-vivo screening and selection. The sgRNA library, WT Cas9 expressing PDX366 cells were resuspended in 150 μL Matrigel ®Growth Factor Reduced Basement Membrane Matrix (Corning, Corning, NY). After the anesthesia, the left flank of the mouse was opened to exteriorize the pancreas, and 8×10^6^ PDX366 cells were injected directly into the pancreas. At this stage, one batch of cells was harvested as “day 0” control sample. For *in vitro* screening, cells were passaged every 3-4 days by 1:3 split with fresh media in 15 cm plates. At least 12 million cells were passaged each time using 3×15 cm plates.

Tumor volumes were monitored by MRI. MRI measurement (University of Virginia Molecular Imaging Core, Charlottesville, VA) was performed after four weeks, at the conclusion of the experiment. Tumors were harvested and weighed, and samples collected for further analysis. Formalin-fixed tumor samples were submitted to the University of Virginia Research Histology Core Lab for processing and H&E staining. Tumor sections were scored by a board-certified pathologist who specializes in gastrointestinal cancers. This study was carried out in strict accordance with the recommendations in the Guide for the Care and Use of Laboratory Animals of the National Institutes of Health. The animal protocol was approved by the Animal Care and Use Committee of the University of Virginia (PHS Assurance #A3245-01).

### Targeted amplification of CRISPR/sgRNA library and sequencing

Tumors from mice and in vitro cultured cells were harvested after four weeks. Entire tumors and all cell pellets were used to obtain genomic DNA. Briefly, tumor samples were minced into small pieces and lysed with 8 ml SDS lysis buffer (100 mM NaCl, 50 mM Tris-Cl pH 8.1, 5 mM EDTA, and 1% wt/vol SDS). Cell pellets were processed in a similar way. Minced tumor samples or cell pellets were treated with 100 μL proteinase K (20 mg/ml) at 55 °C for overnight incubation. The next day, entire lysis solutions were used in EtOH precipitation, and genomic DNA pellets washed with 70% EtOH twice. Pellets were resuspended in RNase-containing water and quantified by Nanodrop. For each DNA sample, 100 μg genomic DNA was used for the first PCR reaction. We ran ten separate PCR reactions with ten μg DNA in a single PCR tube. We used the same outer *Forward Primer* and outer *Reverse Primer* from Sabatini sgRNA library-specific primers for all of the samples (these primers are different from GeCKO Array *For* and *Rev*). Q5-high Fidelity 2X master mix was used as polymerase from NEB (# M0429L). PCR condition for the first PCR was; 98 °C for 30 sec, 18× (98 °C for 10 sec, 63 °C for 10 sec, 72 °C for 25 sec), 72 °C for 2 min. After the first PCR, all reactions were combined (10x 100 μL) in one single Eppendorf tube and vortexed well. For the second PCR, 5 μL PCR reaction mix from the first PCR step was used in 100 μL total PCR reaction. PCR conditions for the second PCR were: 98 °C for 30 sec, 24× (98 °C for 10 sec, 63 °C for 10 sec, 72 °C for 25 sec), 72 °C for 2min. In the second PCR, each sample was amplified with specific forward primers that had a six bp barcode sequence for demultiplexing of our reads during next-generation sequencing and common reverse primer. In this setting, custom sequencing and custom indexing primers for Illumina Sequencing were used. All primer sequences used for library preparation and next-gen sequencing are listed in **Supplemental Table x**. The entire solution from the second PCR was loaded on a 2% gel, and the bands around 270 bp were cut and cleaned with the Qiagen gel extraction kit (a faint band above 270 bp was noticed, likely due to carrying over of primers from the first PCR reaction). Purified PCR products were quantified by using Qubit (Invitrogen), and equimolar amounts of each PCR fragment were mixed and used for subsequent high-throughput sequencing (40 nM DNA in 20 μL). The library was sequenced using the Illumina Miseq platform to get an average of 10 million reads for each sample.

### Data analysis for CRISPR/Cas9 screening

Sequencing reads from CRISPR/Cas9 screenings were first demultiplexed with cutadapt (v. 1.8.3). Sequences of a total length of 56 nt (sequencing barcode and sample barcode) were supplied to the program with the requirements that at least 36 nt of this barcode had to be present in the read, so that it could be assigned to an individual tumor isolated from the PDX model. More than 99% of reads were assigned to one of the three *in vitro* and six *in vivo* samples: cells from the day of injection (further referred to as day 0), control and gem treated *in vitro* samples (one each), and control and gem treated *in vivo* samples (three each). After de-multiplexing and removing sequencing and sample barcodes, the abundance of each sgRNA was assessed and normalized among samples with the use of MAGeCK v. 0.5.2. About 87% of the reads contained correct sgRNA sequences.

Downstream data analysis was performed in RStudio v. 0.99.484 with R v. 3.3.0 following the previous publications(Wang, et al. 2014) with slight modifications. We performed the following analysis to identify potential combinatorial targets of gemcitabine. The first step of this analysis was to calculate the relative abundance of sgRNAs targeting each gene between “day 0” and one of the other eight samples by comparing normalized average counts of all the sgRNAs targeting the particular gene. Since the nongenomic targeting control sgRNAs were well represented in all the samples, they were used to profile the null distribution of Robust Rank Aggregation (RRA) scores when calculating the P values. Based on the negative selection RRA scores, one of the in vivo gem treated samples had a substantially higher sgRNA depletion rate compared to the other two replicates, and thus was excluded from the downstream analysis. Genes consistently depleted in all the samples compared to “day 0” were likely to be essential genes for the PDX cell line, and were removed from the downstream analysis. The second step of the analysis was to calculate log fold change (LFC) of mean read counts between gem treated and control samples for all the retained genes in in vitro and in vivo settings, respectively. In the third step, we ranked all the retained genes based on LFC, and genes significantly depleted (FDR q < 0.1) in both in vitro and in vivo screenings were selected as candidate combinatorial targets of gemcitabine.

### Validation of PRMT5 as a viable CRISPR Screening hit

For validation of PRMT5 after the initial screening, the following sgRNA guiding sequences (sgRNAs) were designed and cloned to generate *PRMT5* knock out cells;

sgRNA1: GGTACCCTTGGTGGCACCAG,
sgRNA2: GGTGATGGCCAGTGTGGATG,
sgRNA3: GTAAGGGGCAGCAGGAAAGC.

Briefly, the oligos that have −5’CACC and −5’AAAC overhangs of the sgRNA guiding sequence were ordered from Eurofins and hybridized to get sticky end double-strand DNA for ligation. The plasmid containing the sgRNA backbone was digested with Bbs.i at 55 °C for 2 hours, followed by CIP treatment at 37 °C for a half-hour. Purified vector backbone from a 2 % gel (60 ng) and hybridized oligos (1 μL from 1-10 nM) was used for the ligation reaction in the presence of T4 ligase.

WT Cas9 and gRNA expressing lentivirus were generated using the HEK293T cell line. mPanc96 and PANC-1 cells were virally infected to express Cas9 and sgRNA to produce stable cell lines. After four days of puromycin selection (2 μg/ml), serial dilution was performed to generate single clones. Once the desired number of clones was obtained, lysates were prepared in RIPA buffer, and Western Blot was performed to determine PRMT5 knockout efficiency.

### MTT Cell Viability

PDX366, mPanc96, and PANC-1 cells were seeded in a flat-bottom 96-well plate (Corning) in triplicate at a density of 1-2×10^3^ cells per well. The following day, cells were treated with gemcitabine (GEMZAR; Eli Lilly) and EPZ015666 (GSK3235025; SelleckChem) or EPZ015938 (GSK3326595; ChemieTek) for 4-5 days prior to MTT (3-(4,5-Dimethylthiazolyl)-2,5-diphenyltetrazolium bromide) to determine effects of drugs on cell viability. Culture media were replaced with fresh RPMI, which had 10% FBS and 10% MTT (5mg/ml) and incubated for 4 hours in a humidified (37 °C, 5% CO2) incubator. 100 μl MTT solvent (10% SDS in 0.01M HCL) was added to each well, and cells were incubated overnight. The absorbance was read at 595 nm.

### Crystal Violet Assay

Pancreatic cancer cells were seeded in a flat-bottom 12-well plate (Corning) at a density of 1-2×10^3^ cells per well. The following day, cells were treated with gemcitabine and EPZ015666 or EPZ015938 for two weeks. Culture media were replaced every week with fresh medium in the presence of drugs. Wells were washed with PBS, then stained for 30 minutes with crystal violet solution (0.4% crystal violet, 10% formaldehyde, 80% methanol). After staining, wells were washed once with PBS and water. The plate was dried out overnight and imaged using a scanner. Colonies were measured and analyzed with ImageJ (National Institutes of Health).

### Annexin V Staining

Annexin V staining was performed to determine the percentage of apoptotic cells. After treatment with gemcitabine and EPZ015666 or EPZ015938, the pancreatic cancer cells were washed with cold PBS, resuspended in Annexin V binding buffer (10mM HEPES, 140mM NaCl, and 2.5mM CaCl2, pH 7.4) with an appropriate amount of FITC-conjugated Annexin V antibody (Life Technologies #A13199), and incubated at room temperature (RT) for 15 minutes. After washing with binding buffer, the cells were resuspended in 2 μg/ml propidium iodide (PI) (Sigma) in PBS plus RNase, incubated at RT for 15 minutes in the dark, and then analyzed using a FACSCalibur flow cytometer (Becton-Dickinson, San Jose, CA, USA).

### BrdU Staining

Pancreatic cancer cells were treated with gemcitabine and EPZ015666 or EPZ015938, incorporated with BrdU (Sigma) for 1 hour, and then fixed by 70% ethanol. BrdU staining was performed according to the manufacturer’s instructions (BD Biosciences, Franklin Lakes, NJ, USA). Briefly, the fixed cells were washed with PBS and then resuspended in 2 N HCl for 20 min to denature the DNA. After washing with 0.1 M Na_2_B_4_O_7_, pH 8.5, to stop acid denaturation, the cells were resuspended and washed with 180 μl 0.5% Tween 20 (Sigma) with 1% normal goat serum (NGS) (Dako, Glostrup, Denmark) in PBS. Then, the cells were incubated with Alexa Fluor 647-conjugated anti-BrdU (mAb) (Invitrogen) for 1 hour at room temperature in the dark. After washing with PBS, the cells were resuspended in 2 μg/ml propidium iodide (PI) (Sigma) in PBS plus RNase, incubated at 37°C for 30min in the dark, and then analyzed by FACSCalibur flow cytometer (Becton-Dickinson, San Jose, CA, USA).

### Western blot

Cells were washed with cold PBS, and then lysed in RIPA buffer (Cell Signaling Technology). After centrifugation at 14,000 rpm for 15 minutes at 4°C, the supernatants were collected, and the protein concentrations were measured using BCA protein assay reagent (BIO-RAD). Subsequently, equal amounts of proteins were separated in NuPAGE 4–12% Bis-Tris gradient gel (Invitrogen #NP0335), and transferred onto nitrocellulose membranes (Invitrogen #B301002). After blocking with 5% milk, the membranes were then probed at 4°C overnight with various primary antibodies: anti-γ-H2AX (Cell Signaling), anti-phospho-Chk1 (Ser345) (Cell Signaling), anti-phospho-Chk2 (Thr68) (Cell Signaling), anti-PRMT5 (Abcam), anti-RPA1 (Abcam), anti-phospho-RPA2 (S4/S8) (Bethyl Laboratories), anti-RPA2 (Ab-2) (Calbiochem), cleaved caspase-3 (Cell Signaling), and anti-β-actin (Sigma), washed with TBST (20 mM Tris, 150 mM NaCl, 0.1% Tween 20; pH 7.6), and incubated with horseradish peroxidase (HRP)-conjugated secondary antibodies (Promega) at room temperature for 1 hour. Finally, after washing with TBST, the antibody-bound membranes were treated with enhanced chemiluminescent western blot detection reagents (GE Healthcare) and visualized with an x-ray film (GE Healthcare).

### Immunofluorescence staining

Cells grown on glass coverslips (VWR) were rinsed with PBS, and then fixed in 4% formaldehyde for 15 minutes. The cells were subsequently treated with 0.2% Triton X-100 in PBS for 10 minutes. After blocking with 2% BSA in PBS containing 5% FBS at RT for 30 minutes, cells were incubated with an appropriate primary antibody γ-H2AX (Cell Signaling) for 2 hours. Then the cells were washed with PBS and incubated for 1 hour with secondary antibody [Alexa Fluor-488 goat anti-mouse immunoglobulin G (IgG) (H+L) conjugate or anti-rabbit IgG (H+L) conjugate (Invitrogen)]. After washing with PBS, the coverslips were dried, and then reversely covered onto slides (Fisher Scientific) by adding mounting medium with 4′,6-diamidino-2-phenylindole dihydrochloride (DAPI) (Vector Laboratories). A LSM-710 confocal microscope (Zeiss) was used to obtain fluorescence images.

### Comet assay

The comet assay measures DNA damage in individual cells. It was performed according to the instructions of the OxiSelect Comet Assay Kit (Cell Biolabs). Briefly, microscope slides were first covered with a normal melting point agarose to create a base layer. Then, 1 - 2 ×10^5^ of cells were embedded into 75 μL of low-melting-point agarose at 37°C, and the gel was cast over the first agarose layer. Then slides were immersed into a lysis buffer and kept for 1 hour at 4°C. After cell lysis, the slides were electrophoresed in alkaline electrophoresis buffer (300 mM NaOH, one mM EDTA, pH13). The slides were then stained with Vista Green DNA dye. Comet tails were measured using Image J.

### NHEJ and HR repair assays

NHEJ and HR assays to examine the repair efficiency of I-SceI inducible-double strand breaks (DSBs) were performed using NHEJ/DsRed293B(Golding, et al. 2009) and HeLa DR13-9(Ransburgh, et al. 2010) cell lines, respectively, as previously described with slight modifications (Lee, et al. 2017). Briefly, 3 × 10^5^ cells were plated on 6 well dishes, and 2 μg I-SceI expression vector pCβASce was transfected using Lipofectamine2000 (Invitrogen). After 24 hr post-transfection, the indicated amount of gemcitabine and/or EPZ015666 was incubated with cells for 24 hr. The DsRed- and GFP-expressing cells were counted in flow cytometric analysis (BD FACS Calibur and CellQuest Pro) by the FL2 and FL1 channels for NHEJ and HR repair efficiency, respectively. The % of fluorescent positive cells in the treatment of gemcitabine and/or EPZ015666 was normalized to that of the non-treatment cells (Ctrl) transfected with pCβASce to calculate the relative repair efficiency.

### RPA overexpression

WT RPA1/2/3 and GFP were overexpressed in the PRMT5 KO cell line by co-transfection of WT RPA1/2/3 (Addgene) and pCMV-GFP plasmid with 5:1 ratio. The wild-type cell line was transfected with pCMV-GFP plasmid alone as a control. 10 μg total DNA was used in the presence of 30 μl of Fugene6 reagent in 10 cm plate dish that had 70 % confluency. After transfection 24 hours, GFP positive cells were sorted by a FACS Aria cell sorter.

### *In vivo* xenograft experiments

All animal care and experimental procedures were carried out in accordance with protocols approved by the University of Virginia School of Medicine Animal Care and Use Committee (PHS Assurance #A3245-01). To develop xenograft tumors, Control sgRNA infected WT cells and PRMT5-KO cells were subcutaneously injected into the dorsal flanks of 8 week-old nude mice, which were obtained from the Jackson Laboratory (Bar Harbor, ME, USA). When the tumors were visible (approximately 30 mm^3^ in volume), the mice received respective gemcitabine treatments via intraperitoneal (i.p.) injection. After weekly monitoring, time to appearance of the tumor was recorded, and the tumor volume was measured by caliper. The tumor volume was calculated as follows: volume = longest tumor diameter × (shortest tumor diameter)^2^/2. After 35 days of treatment, the mice were euthanized by CO2 inhalation, and the tumor tissues were collected for further analyses.

### RNA-Seq and library preparation

Control sgRNA (CgRNA) and PRMT5-KO cells were treated with gemcitabine (200 nM) or EPZ015666 (500 nM) for 48 hours. Total RNA was purified using the RNeasy mini kit (Qiagen #74104) by following the kit instructions. mRNA was isolated by using NEBNext Poly(A) mRNA Magnetic Isolation Module (New England Biolabs # 7490S). RNA-Seq libraries were prepared using the NEBNext Ultra Directional RNA Library Prep Kit for Illumina (New England Biolabs # E7420S) by following the company’s protocol. A Qubit measurement and bioanalyzer were used to determine the library quality.

### Data analysis for RNA-Seq

General sequencing data quality was examined using FastQC (v. 0.11.5). RNA-Seq data were aligned to the human reference genome (hg19) using HISAT2 (Kim, et al. 2015) (v. 2.1.0) with the default paired-end mode settings. The resulting sam files were sorted by reading names and converted to bam files using samtools(Li, et al. 2009) (v. 1.9) sort command. For ATAC-Seq, sequencing reads mapped to mitochondria DNA were removed from the bam files using the samtools view. Then the bam files were sorted by mapping position and indexed using corresponding samtools commands. The sorted and indexed bam files were first converted to bigwig files for visualization in the UCSC Genome Browser (https://genome.ucsc.edu/) to avoid technical alignment errors. Next, the bam files were quantified against gencode (v27lift37) annotation using Stringtie(Pertea, et al. 2016) (v. 1.3.4d) with the default settings.

After obtaining the gene count matrix from Stringtie, we imported it into R and normalized the data following the pipeline of DESeq2 (Love, et al. 2014). Specifically, to ensure a roughly equal distribution for all the genes across samples, we used rlog transformation to stabilize expression variance for genes with different expression levels. Then samples were clustered according to Euclidean/Poisson distances to make sure replicates are clustered together. By calling the DESeq function, we determined genes with significant expression changes between the *PRMT5* WT and KO samples thresholding at an adjusted P value of 0.01. Heatmaps were produced using the pheatmap R package. All other plots were generated using ggplot2. Gene set enrichment analysis(Subramanian, et al. 2005) were performed using the GSEA website (http://software.broadinstitute.org/gsea/index.jsp) and the stand-alone GSEA program referencing the Molecular Signatures Database (MSigDB).

### Data analysis for three publicly available PDAC studies from GEO and TCGA

Data were downloaded from GEO (https://www.ncbi.nlm.nih.gov/geo/) for studies GSE28735, GSE16515, and GSE93326. Normalized PRMT5 expression was compared between relevant sample groups using appropriate student's t-test. Analyses were performed, and plots were generated in RStudio v. 0.99.484 with R v. 3.3.0. The survival analysis for PDAC patients with high (> 0.5 standard deviation [s.d.]) and low expression (< 0.5 s.d.) of PRMT5 was carried out through cBioPortal (www.cbioportal.org).

## Acknowledgment

The study was initially funded by a pilot project award (to Dr. Adli and Dr. Bauer) from the University of Virginia Cancer Center. Additional resources from Pinn Scholar Award (to Dr. Adli) and from NIH/NCI 1R01 CA211648-01 award were mobilized to complete the work. We are thankful to all the members of Adli and Bauer labs for critical discussions during this study.

## AUTHOR CONTRIBUTION

MA conceptualized the study, supervised the experiments, and wrote the manuscript. XW performed the validation and mechanistic experiments. JY performed the computational analysis. CK performed the initial screenings. KYL and AD helped with the DNA rapir assays. SA, PA, SL and XW in consultation with TWB, performed *in vivo* experiments. YMD, AHH, and EY helped with *in vitro* experiments.

