## Supplementary Figures for "Targeted CRISPR screening identifies PRMT5 as synthetic lethality combinatorial target with gemcitabine in pancreatic cancer cells"

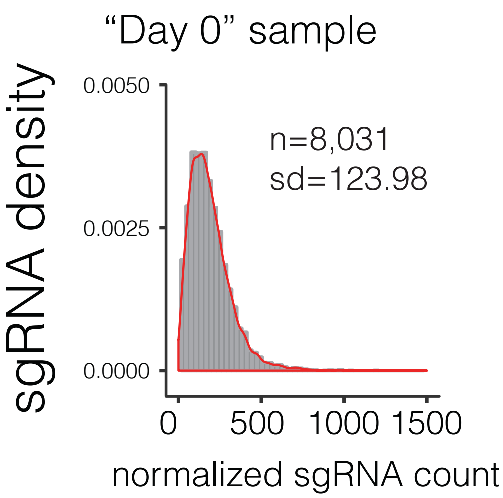


**Supplementary Figure 1:** The density plot shows sgRNA read count distributions in Day 0 sample.

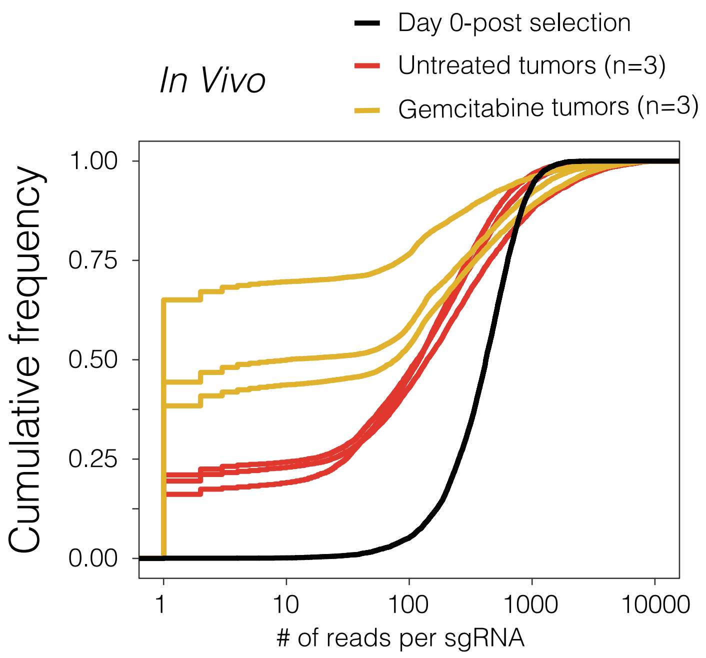


**Supplementary Figure 2:** The cumulative frequency plot shows the fraction of sgRNAs with indicated number of reads detected in Day 0, Untreated tumors and Gem treated tumors.

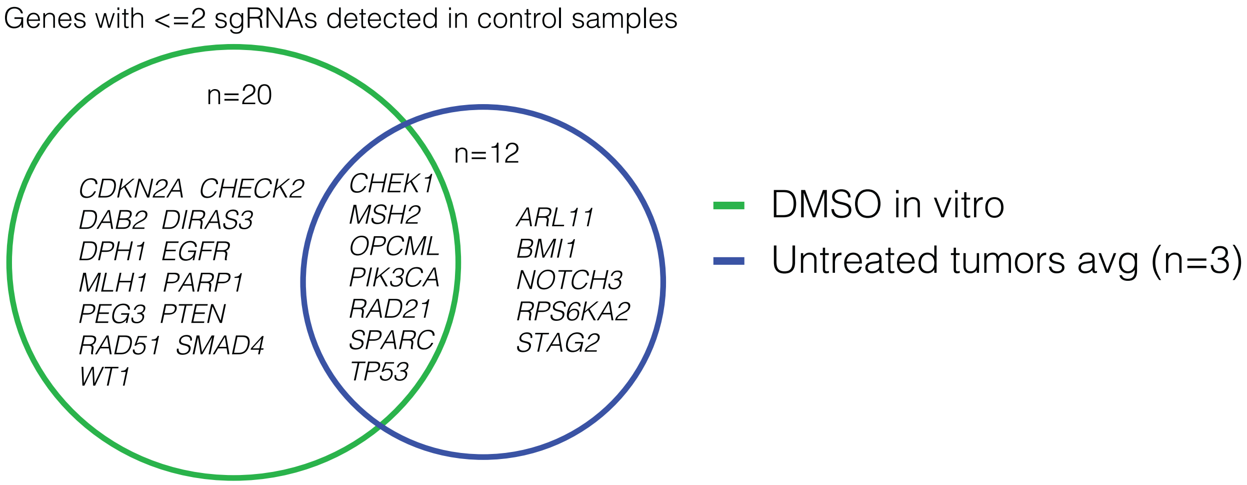


**Supplementary Figure 3:** The Venn diagram shows the genes which were represented by 2 or less sgRNAs *in vitro* and *in vivo*, indicating that their depletion was created significant lethality in all samples.

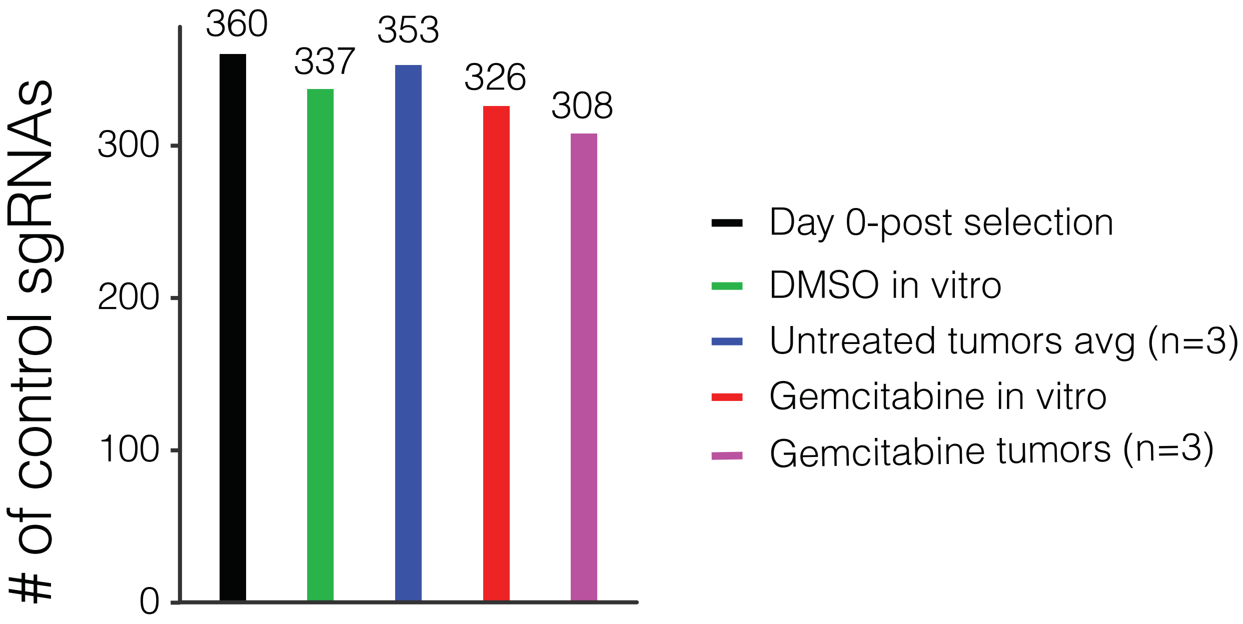


**Supplementary Figure 4:** The bar plot shows number of control sgRNAs (total # in the library: 360) detected in each of the indicated samples.

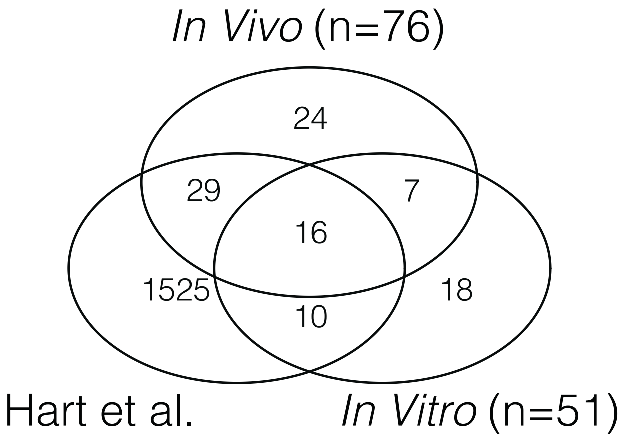


**Supplementary Figure 5:** The Venn diagram shows number of fitness genes detected in *in vivo* and *in vivo* and their comparison with the previously identified "core fitness" genes through genome wide screening by *Hart et al.*[1]

1 Hart, T. *et al.* High-Resolution CRISPR Screens Reveal Fitness Genes and Genotype-Specific Cancer Liabilities. *Cell* **163**, 1515-1526, doi:10.1016/j.cell.2015.11.015 (2015).
